# Mouse Auditory Cortex Undergoes Asynchronous Maturation in the Right and Left Hemispheres

**DOI:** 10.1101/2024.01.16.575905

**Authors:** Ashlan Reid, Demetrios Neophytou, Robert Levy, Hysell V. Oviedo

## Abstract

Despite the significance of lateralized auditory processing in human cognition, there are limited studies in animal models exploring the developmental mechanisms of this cortical specialization. Here, we find that cellular and network signs of maturity in the Auditory Cortex (ACx) appear earlier in the right hemisphere in male mice. We further demonstrate that persistent, experience dependent map reorganization is confined to the hemisphere that is actively maturing and can be differentially engaged by temporally limited manipulations of the sensory environment. Our data suggests that differential timing in hemisphere development could lead to lateralized auditory functioning.

## Main

Sensory “critical periods” are brief developmental phases when long-term circuit structure is shaped by neural activity reflecting sensory information in the current environment (Fig. 1a)^1^. Sound exposure during the auditory critical period selectively shapes Auditory Cortex (ACx) representations, whether for experimentally controlled tone pips or for the cadence of a child’s household language(s) ^2-4^. In humans and mice, a fundamental feature of mature auditory sensory processing is the allocation of specialized cognitive functions to the Left and Right ACx ^5-7^. Failure of the ACx to develop lateralized function is a common endophenotype of human cognitive disorders, such as Autism Spectrum Disorders and Schizophrenia ^8,9^. In rodents, selective deactivation of one of the auditory cortices impairs distinct auditory processing functions^10-12^. In children, the development of functionally lateralized auditory evoked potentials occurs during the first 3 years of life ^13, 14^, and abnormal auditory experiences during infancy lead to degraded language abilities^15^. Altering the acoustic environment during development also results in impaired spectral tuning and abnormal circuit patterns in rodents^16,6,17^. Although the importance of lateralized auditory processing in human cognition is well recognized, there are few studies in animal models investigating the developmental mechanisms of this lateralization. Here, we set out to compare mouse Left and Right ACx functionality across developmental time windows. We identify time periods when Left and Right ACx are maturing asynchronously, and manipulation of the sensory environment during this time leads to lasting asymmetries in adult tone representations.

**Fig. 1:**
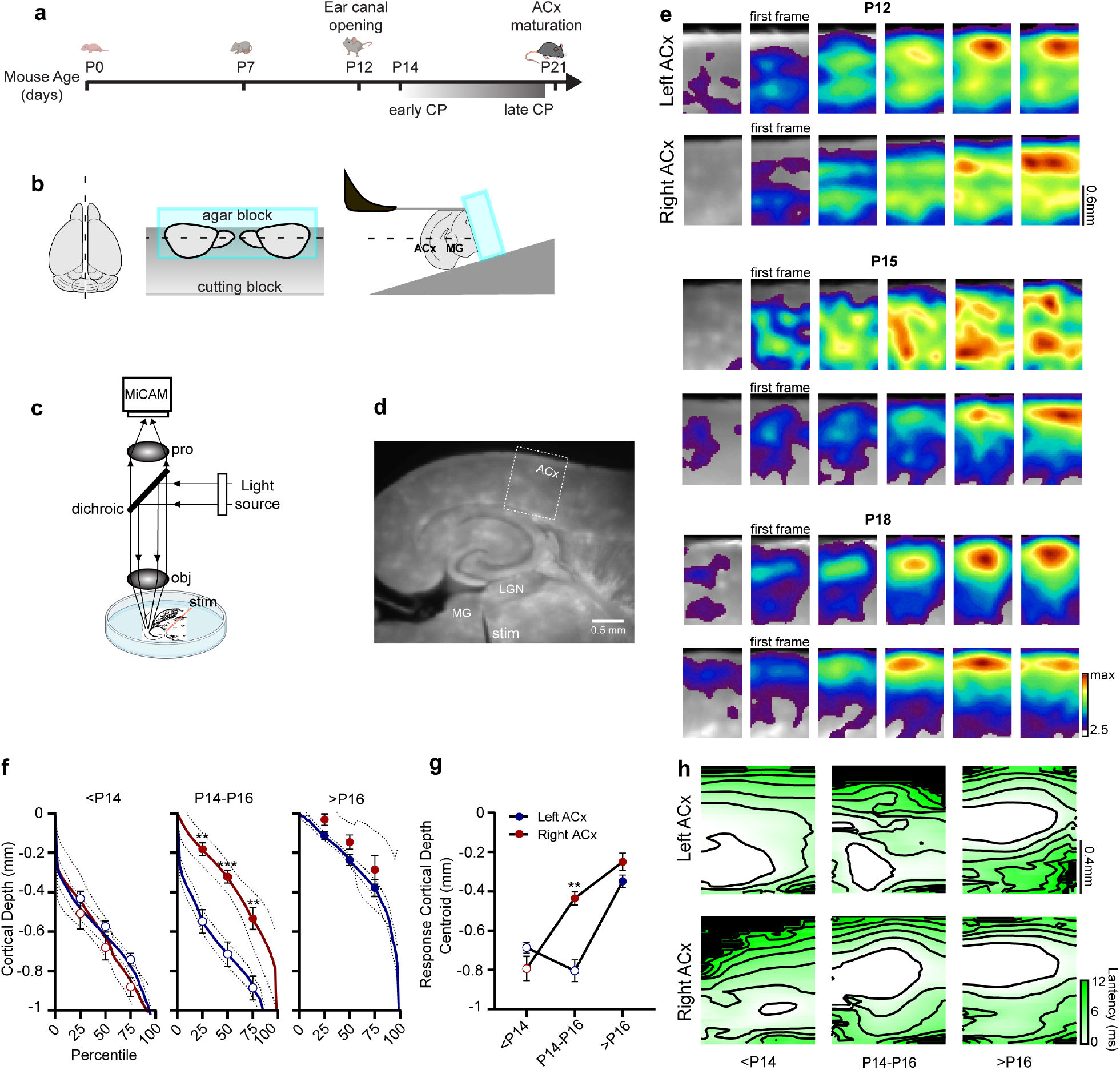
Hemispheric differences in the maturation of thalamocortical input to the ACx. **a**, timeline of major milestones in the development of the auditory system and ACx tone critical period. **b**, connected bilateral TC slice preparation developed to study the maturation of the Left and Right ACx in the same animal. **c**, experimental set-up for VSD. **d**, picture of a connected TC slice stimulated in the MGBv with ACx labeled where voltage changes were measured. **e**, movie frames (4ms rate) for Left and Right ACx TC responses; *first frame*: first movie frame determined to show a significant TC response (see Methods). **f**, cumulative binned depths of responsive locations for pixels in the upper quartile (>75th percentile) of signal magnitude for the *first frame*. Left ACx: *blue*, Right ACx: *red*; mean: *solid lines*, 95% confidence intervals: *dotted lines*. Circles indicate 25th, 50th (median), and 75th percentiles of the population average and S.E.M. There was no significant difference in the youngest group <P14 (*empty circles*, Left panel, unpaired Welch’s t-test, Holm-Šídák method of correction for multiple comparisons; n=9 and 7 slices for Left and Right ACx, respectively). A significant difference between the Left and Right ACx was observed at P14-16 (*filled circles*, middle panel: 25th percentile p=0.00107, 50th percentile p=0.000939, 75th percentile p=0.0041; n=7 and 6, unpaired Welch’s t-test, Holm-Šídák correction). There was no significant difference in the oldest group (>P16, right panel, n=9 and 5; unpaired Welch’s t-test, Holm-Šídák correction). See Supplemental table 1 for within hemisphere statistical tests for these data. **g**, Centroid of response calculated for the first frame across age and hemisphere groups. The centroid was not significantly different for ages <P14 nor >P16 (unpaired Welch’s t-test, Holm-Šídák correction; for <P14 n=9 and 7; for >P16 n=9 and 5). The only significant difference in centroid response between the Left and Right ACx occurred at ages P14-16 (p=0.000747, n=7 and 6; unpaired Welch’s t-test, Holm-Šídák correction). **h**, Population averaged contour plots reporting the spatial distribution of thalamic input response latency.

Crucial insights into mesoscale dynamics of developing and mature auditory cortical circuits have been revealed by physiological assays performed *in vitro* using connected auditory thalamocortical (TC) slice preparations^18-20^. As described, this preparation involves the loss of one hemisphere, usually the right, to achieve the correct compound slice angle. To compare Left and Right ACx circuits in the same animal, we developed a new preparation technique for simultaneous retrieval of connected TC slices from both hemispheres (Fig. 1b, see Methods). Directly comparable to previous studies, we record optical signals (Fig. 1c) in voltage-sensitive-dye stained slices (Fig. 1d) to capture the full range of sub- and suprathreshold neuronal depolarizations driven in the ACx by electrical activation of thalamocortical (TC) axons.

As expected, auditory thalamocortical responses in both Left and Right ACx changed significantly with maturity of the animal (Fig. 1e-h). Consistent with the standing model of immature TC responses initiating in the cortical subplate^21,19,22,23^, we found that in ages <P12, the Left ACx TC response arose first in lower layers and showed delayed activation of layers 2/3-4 (Fig 1e-h). By contrast, in mature (>P16) Left ACx, the thalamocortical response is initiated in layers 4 and lower 2/3, indicating a developmental change in circuit architecture. We next determined that this shift in TC response location also occurs during development in the Right ACx (Fig 1e-h). Surprisingly, in age- and animal-matched slices, this significant laminar shift arose in the Right ACx at earlier ages compared to the Left ACx (P14-P16, Fig1e-h). Consequently, the differences in thalamocortical response were significant at ages P14-P16, with the Right ACx displaying signs of maturity and the Left ACx appearing less mature (Fig 1f-g).

Previous studies have shown that relative response latency within layers is another metric to indicate the spatial density of direct thalamic input^19,24^. In population averaged contour plots reporting the spatial distribution of response latencies, the region of earliest response was spatially shifted to upper layers in the mature ACx (Fig 1h, right column). Again, at ages P14-P16 this shift is apparent in the Right ACx and not yet present in the Left ACx. Across age- and animal-matched studies, both spatial and temporal data support earlier ACx maturation in the right hemisphere.

If structural changes in thalamocortical input suggest asynchronous maturation between Left and Right ACx, we reasoned that other mature circuit phenomena could also emerge in the two hemispheres at different ages. Synaptic contacts between excitatory and inhibitory neurons undergo developmental shifts^25,26^, and maturation of inhibitory neurons is a crucial trigger of activity-dependent circuit refinement during the critical period^27,1^. We therefore measured changes in spontaneous inhibitory postsynaptic currents (sIPSCs) across ages in pyramidal cells in L4 of age- and animal-matched Left and Right ACx (Fig 2a, see Methods).

**Fig. 2:**
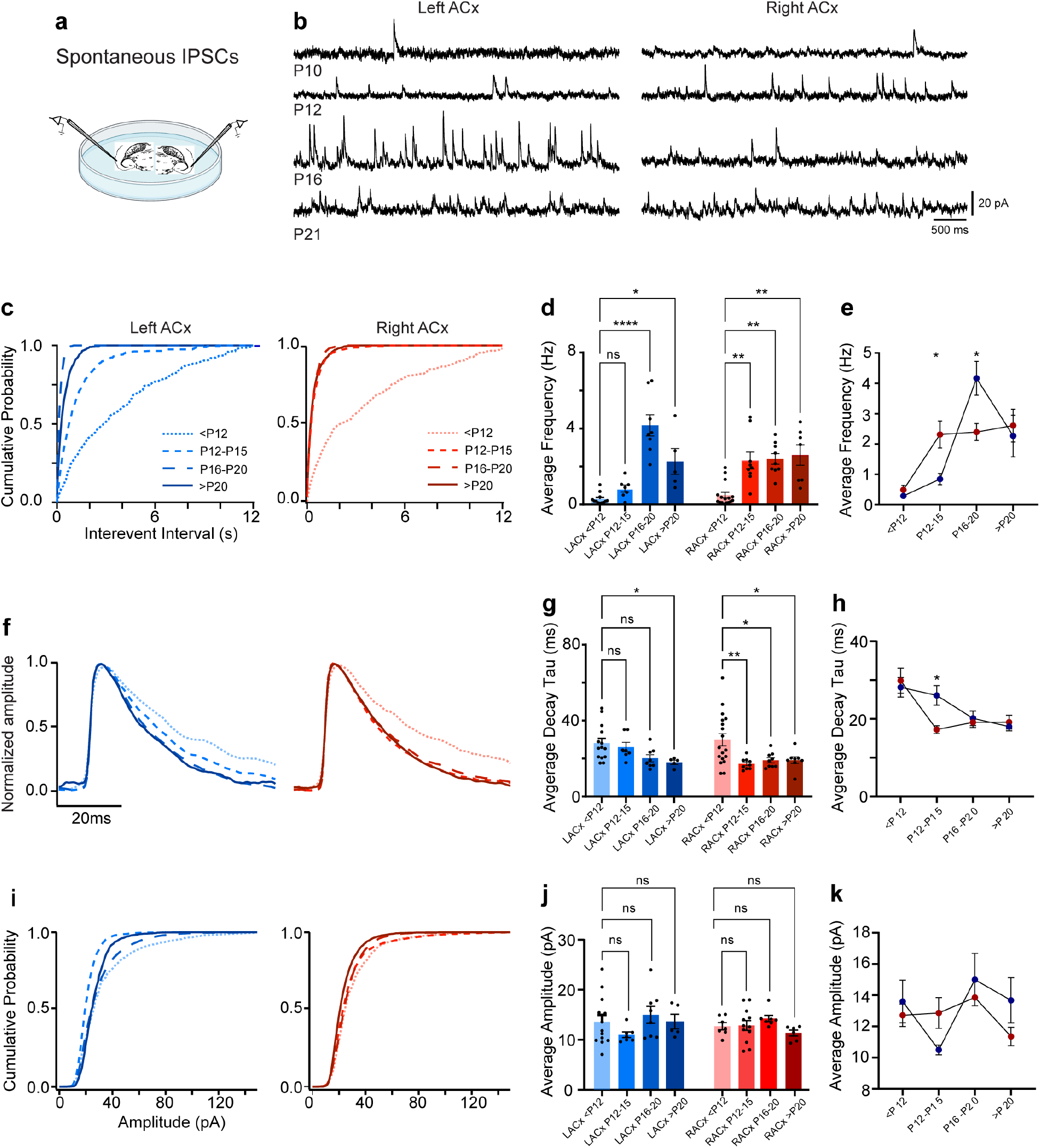
Hemispheric differences in the maturation of inhibitory synaptic input in the ACx. **a**, Slices were collected from both hemispheres for age-matched and within-animal comparison of sIPSCs. **b**, Voltage clamp recordings were performed sequentially from excitatory neurons in L4 of the Left (left traces) and Right (right traces) ACx (order was randomized between animals). Sample traces of sIPSCs recorded at four ages: starting from prior to ear canal opening (P10) to end of tone critical period (P21). **c**, Cumulative histograms of sIPSC interevent interval from the Left and Right ACx at 4 age groups. **d**, Quantification of developmental changes in mean sIPSC frequency within each hemisphere. (Comparison to youngest group within each hemisphere. Left ACx: <P12 (n=12 cells) vs. P12-P15 (n=7), p=ns; <P12 vs. P16-P20 (n=8), p=<0.0001; <P12 vs. >P20 (n=5), p=0.0236. Right ACx: <P12 (n=15) vs. P12-P15 (n=7), p=0.0038; <P12 vs. P16-P20 (n=9), p=0.0015; <P12 vs. >P20 (p=7), p=0.0055, Kruskal-Wallis, post hoc Dunn’s multiple comparisons test). **e**, Comparison of developmental changes in mean sIPSC frequency between the hemispheres (<P12, p=ns, n=12 and 15 cells, Left and Right ACx, respectively; P12-P15, p=0.031099, n=7 and 7; P16-P20, p=0.032951, n=8 and 9; >P20, p=ns, n=5 and 7, Multiple Mann Whitney test, Holm-Šídák correction). **f**, Normalized sIPSCs illustrate developmental changes in the decay time constant. **g**, Quantification of sIPSC decay time constant maturation within each hemisphere. (Comparison to youngest group within each hemisphere. Left ACx: <P12 (n=14) vs. P12-P15 (n=5), p=ns; <P12 vs. P16-P20 (n=8), p=ns; <P12 vs. >P20 (n=5), p=0.0109. Right ACx: <P12 (n=18) vs. P12-P15 (n=9), p=0.0076; <P12 vs. P16-P20 (p=9), p=0.0318; <P12 vs. >P20 (n=6), p=0.0277, Welch’s ANOVA test, post hoc Dunnett’s t-test). **h**, Comparison of developmental changes in sIPSC decay time constant between the hemispheres (<P12, p=ns, n=14 and 18; P12-P15, p=0.04821, n=5 and 9; P16-P20, p=ns, n=8 and 9; >P20, p=ns, n=5 and 6 unpaired Welch’s t-test, Holm-Šídák correction). **i-k**, Quantification, and comparison of sIPSC amplitude during development within hemispheres shows no statistically significant difference (Comparison to youngest group within each hemisphere. Left ACx: <P12 (n=13 cells) vs. P12-P15 (n=7), p=ns; <P12 vs. P16-P20 (n=8), p=ns; <P12 vs. >P20 (n=5), p=ns. Right ACx: <P12 (n=17) vs. P12-P15 (n=8), p=ns; <P12 vs. P16-P20 (n=9), p=ns; <P12 vs. >P20 (p=6), p=ns, Welch’s ANOVA test, post hoc Dunnett’s t test). Comparison of sIPSC amplitude during development between hemispheres shows no statistically significant difference (<P12, p=ns, n=13 and 17 cells, Left and Right ACx, respectively; P12-P15, p=ns, n=7 and 8; P16-P20, p=ns, n=8 and 9; >P20, p=ns, n=5 and 6; Unpaired Welch’s t-test, Holm-Šídák correction).

To assess maturation of cortical inhibitory tone, we analyzed key sIPSC properties known to change developmentally^28,26,29^. First, we observed that the frequency of sIPSCs increases significantly at P12-15 in the Right ACx but not the Left ACx (Fig 2b-c). Moreover, the average frequency of sIPSCs remains stable in the Right ACx with no further changes beyond the P12-15 time window (Fig 2b-d). Interestingly, the Left ACx shows a more dynamic change in sIPSC frequency during development. At P16-20, the Left ACx showed a brief increase in average frequency and shortened interevent intervals (IEIs, Fig 2b-d), consistent with prior findings of transient hyperconnectivity in its superficial layers^30^. We did not observe the transient sIPSC frequency spike in the Right ACx, suggesting that absolute differences may exist between the maturation pathways of the two auditory cortices. After P20, sIPSC average frequencies and IEI distributions in the Left ACx are comparable to the same measures in mature Right ACx (Fig 2e).

Next, we examined the decay time constant of sIPSCs, which has been shown to decrease as intracortical inhibition matures^31,32,26^. As expected, we found that the decay time shortens with age in both auditory cortices (Fig 2f). However, the Right ACx demonstrated a shift to shorter decay times at P12-15 and no further changes with age (Fig 2f-h). By contrast, in the Left ACx the age-related decrease was less abrupt and did not reach significance until later ages (Fig 2g). These observations may reflect lateralized changes in GABA_A_ receptor kinetics during development, potentially due to changes in receptor subunit composition^28^. Finally, we found sIPSC amplitude to be stable across ages and hemispheres, with no significant trends between age groups (Fig 2i-k).

Developmental plasticity influenced by experience leads to the formation of finely-tuned representations of the external world in the mature brain. To determine if the asynchronous maturational events in the ACx have an impact on experience-dependent plasticity, we tone-reared mouse pups with patterned 7kHz pure tone pips^19,17^ from P12-15. During this time, our in vitro results predict that the Right ACx is transiently more sensitive to the sensory environment compared to the Left ACx (Fig 3a). After the tone-rearing period, mice were returned to the colony until adulthood. To measure the impact of this juvenile transient tone exposure on mature tonotopic representations, we performed *in vivo*, anesthetized, bilateral extracellular recordings using multichannel silicon probes in adult mice from control and transiently tone-reared cohorts. Spike times were determined blind to tone stimuli presentation and were grouped into clusters based on spatiotemporal template matching using Kilosort. Clusters were included in determining the tone response properties of a given location based on the presence of spikes time-locked to tone presentation (see Methods).

**Fig. 3:**
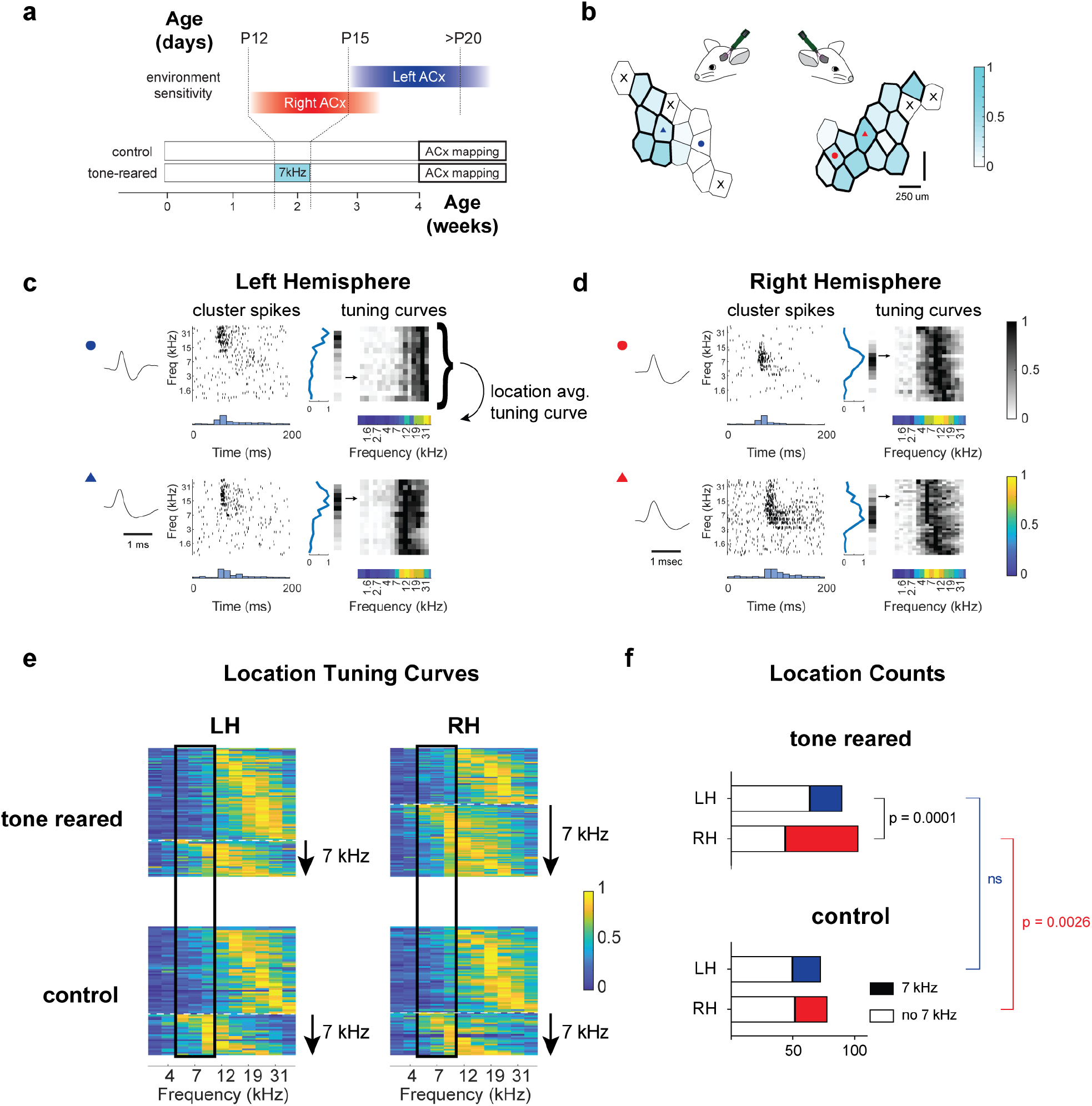
Experience-dependent map reorganization is confined to the hemisphere that is actively maturing. **a**, mouse pups were either reared in control conditions (without exposure to acoustic manipulation), or exposed to 7kHz tone pips between P12-P15. **b**, Between P33-P56 bilateral extracellular recordings from the Left and Right ACx were performed in anesthetized mice from both control and tone-reared groups using 32 channel, dual-shank silicon probes. Bilateral tessellation maps from one animal showing recording locations and colored according to the fraction of spike clusters at that location responsive to 7kHz. **c-d**, Examples of putative individual neuron tone responses (locations indicated by circles and triangles in tessellation maps) for the Left and Right ACx, respectively. Raster plots, peristimulus time histograms, and normalized tuning curves are shown for each neuron. Grayscale heatmaps show normalized tuning curves for all clusters at the same location; arrow points to the row containing the tuning curve of the example. Colored heatmap represents the average normalized tuning curve calculated for that location and used for further analysis. **e**, Recording location tuning curves from control and tone-reared mice in the Left and Right ACx, sorted by the tuning curve maxima and divided into areas not responsive to 7kHz (above the dotted line) and responsive to 7kHz (below the dotted line). Vertical rectangle shows the location of 7kHz +/-⅓ octave on the frequency axis. **f**, Population counts of recording locations responsive (solid) and not responsive (empty) to tones close to 7kHz. Results from Fisher’s Exact Test showing that the Right ACx had a significantly larger area responsive to tones close to 7kHz compared to the Left (number of recording locations: Left and Right ACx tone reared n=90 and 105 total recording locations from 9 and 10 mice, respectively; Left and Right ACx control n=73 and 78 total recording locations from 11 and 12 mice, respectively).

Confirming our novel hypothesis, in adult mice briefly exposed to 7kHz tones as juveniles, the Right ACx was significantly more responsive to tones close to 7kHz—within a third of an octave—when compared to the Left ACx (Fig 3b-d). Furthermore, comparing proportions of 7kHz responsive cortical locations between control and tone-reared cohorts showed significant trends reflecting over-representation of 7kHz in the Right ACx but not in the Left ACx (Fig 3e-f). These *in vivo* results confirm that brief tone-rearing during development differentially influenced the adult tonotopic maps in the Right and Left ACx. This apparent hemisphere-specific, experience-dependent reorganization coincided with the developmental trends we observed *in vitro*. Together, our data indicate a temporal window in which Right ACx is more sensitive to manipulation of the sensory environment, providing an opportunity for differential representations to emerge.

Network, intracellular, and plasticity data suggest earlier Right ACx maturation in male mice, reminiscent of earlier right hemisphere maturation observed in the human auditory system^14^. Given the abundance of changes taking place in early postnatal development, including those physically intrinsic to the animal (e.g., ear canal opening) and in the external environment (e.g., littermate vocalizations), a 2-4 day difference in the critical period time window between the auditory cortices could dramatically influence the nature of the acoustic inputs co-occurring with the molecular events driving circuit maturation. Thus, a hemisphere-specific temporal shift in ACx maturational trajectory has the potential to precipitate the lateralized functionality found in adult cortical circuits and disrupted in various brain disorders^9,8^. Our findings support the utility of the mouse as an animal model to dissect the mechanistic underpinnings of development leading to functional specialization for communication processing in the Left cerebral hemisphere. The delayed and extended maturation of the Left ACx may facilitate the brain’s ability to fine-tune circuits for spectrotemporal sensitivity, crucial for recognizing the statistical structure of species-specific vocalizations. Importantly, delayed maturation may also render the Left ACx more vulnerable to injury^33^. Lastly, our findings advise caution for experimental designs in which developing partner hemispheres are assumed to be identical and therefore suitable for providing control data. Further studies are needed to test whether the divergent maturational trajectories we observed causally lead to hemispheric specializations in healthy circuit structure and function.

## Methods

Experiments were performed using male CBA/J mice in strict accordance with the National Institutes of Health guidelines, as approved by The City College of New York Institutional Animal Care and Use Committee. For *in vitro* studies, male mice aged P8-P25 were anesthetized with 4% isoflurane and then decapitated. Brains were removed and placed into chilled carbogen-bubbled cutting solution composed of (in mM): 110 choline chloride, 25 NaHCO3, 25 d-glucose, 11.6 sodium ascorbate, 7 MgCl2, 3.1 sodium pyruvate, 2.5 KCl, 1.25 NaH2PO4, and 0.5 CaCl2. In experiments where thalamocortical connectivity was not required, slices were cut along the horizontal plane on a Leica vibratome, using standard in vitro slice preparation procedures with cyanoacrylate glue and a flat stage. The tissue blocking approach to retrieve connected thalamocortical slices from both hemispheres is depicted in Fig1b. First, the brain was hemisected along the midline to expose a surface of each hemisphere along the sagittal plane. The cut surfaces were then affixed with cyanoacrylate glue to a rectangular block of ∼3% low melt agarose as shown. The agar block and two brain hemispheres were glued to a 15-degree wedge, which was printed out of Nylon 12 by Shapeways (Livonia, MI, USA).

Once slices were obtained, they were transferred to artificial cerebrospinal fluid (ACSF) containing (in mM): 127 NaCl, 25 NaHCO3, 25 D-glucose, 2.5 KCl, 1 MgCl2, 2 CaCl2, and 1.25 NaH2PO4 and continuously bubbled with carbogen. Slices were incubated in a recirculating chamber filled with ACSF warmed to 32°C for one hour and then held at room temperature for the duration of the experiment. For voltage-sensitive dye (VSD) imaging preparations, slices were individually stained for 40 to 90 minutes in a miniaturized recirculating bath chamber filled with 15mL total of room temperature, carbogen-bubbled ACSF with the addition of 15uL of 5mg/mL Di-4-ANNEPS (Thermofisher #D1199) in high purity ethyl alcohol (EtOh), so that the final concentration of dye in the chamber was 5ug/mL and the final concentration of EtOH was 0.1% by volume. The stained slice was then placed in a large volume (∼300mL) recirculating, carbogen-bubbled, room temperature ACSF incubation chamber for a minimum of 20 minutes to remove excess unbound dye and any particulate accumulation before imaging. For VSD optical signal acquisition, individual slices were transferred to a room temperature submersion-style recording chamber mounted on a modified upright microscope (BX51-WI; Olympus). Optical recordings were obtained as single trial, 4ms frame rate, 1028 ms total length, movies using a CCD camera (MiCam02, Brainvision) and corresponding supporting hardware and software from BrainVision. Illumination was delivered using a halogen lamp (MHAB-150W, Moritex) and a dichroic filter cube was custom designed to maximize the excitation, collection, and rejection of the appropriate optical spectra (excitation Edmund #86-354, emission Edmund #84-745, dichroic Semrock FF560-Di01-25x36). Light was delivered and collected via a 4× objective (NA, 0.28; Olympus) and passed through a 0.25× demagnification step (U-TVO.25XC; Olympus) before reaching the camera, resulting in measured pixel dimensions of approximately 31μm x 36 μm. Timing of the lamp shutter and electrical stimulus delivery were precisely controlled by the Brainvision camera system. Electrical stimuli consisted of single 100us pulses of constant current delivered to the thalamocortical axon bundle, medial in the slice with respect to the rostral tip of the hippocampus, using an AMPI stimulus isolation unit triggered by an FHC Pulse-01 and single pole tungsten electrodes modified to be 50-200 kOhm in resistance.

For VSD image processing, we combined and adapted procedures reported in previous studies^24,19,34^. Between 2 and 6 trials at a given stimulus electrode location were collected as time-series movies, averaged frame-by-frame, and then smoothed with a 3x3-pixel flat filter. To align slices across experiments and account for non-standard camera pixel arrangement, movie frames were linearly interpolated to a grid of locations spaced evenly 25 um in each direction and rotated so that the vertical axis of the grid was perpendicular to the layered organization of the cortex in the region of the ACx. Frames were then individually smoothed with a 3x3-pixel gaussian filter and cropped to a rectangle of 1400um in the vertical dimension (across cortical layers) and 800um in the horizontal (anterior-posterior). There was no signal conditioning in time. In voltage-sensitive dye signals of this nature, an increase in cellular membrane voltage is observed as a decrease in raw signal amplitude. Therefore, we invert the signal polarity such that membrane depolarization is reflected as an increase in the optical signal. We present the stimulation elicited change in fluorescence (dF/F) in terms of z-score, or number of standard deviations above the baseline subtracted mean, to account for differences in technical and biological variability in the slice preparations.

To describe and compare the spatiotemporal dynamics of the cortical response in a movie, we first determined the movie frame demonstrating the earliest indication of a significant response after the stimulus (termed “first frame” in Fig 1e). To make the determination of the first frame, a threshold was applied to the image (60 x 80 pixels) and the number of non-contiguous pixels (each 25um x 25um) with a value higher than that threshold were counted. A table was generated for each movie wherein the first frame was calculated for a range of threshold values (z-score 2 to 7.5 in steps of 0.5) and area values (4 to 84 pixels in steps of 4 pixels). The most commonly occurring value (mode) in each table was selected to be the first frame for that movie. All 43 first frames occurred within 3 frames (8 to 16 ms post-stimulus latency) and did not show a trend between experimental groups.

To assess the spatial location of the initial response over the cortical layers, we binarized the first frame for each movie with a z-score threshold set to the 75% signal level and binned the depths of the above-threshold pixel locations into cumulative histograms starting at the cortical surface. This cumulative depth histogram approach (Fig 1f) aids both interpretation and statistical testing of the distribution of the response across the population, since the cortical depths containing specific percentiles of the total responding areas are represented. Across the age and hemisphere groups, the average depth histograms are reported along with 95% confidence intervals (Fig 1f solid and dashed lines, respectively). In addition, we indicate mean and S.E.M. of the 25th, 50th (median), and 75th percentiles in Fig 1f (circles).

For a single value to quantify the mean response location across cortical layers, we calculated the vertical location of the centroid of the first frame image using the equation:

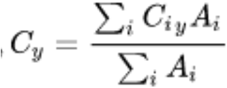

We further sought to quantify the response across time as an indication of direct functional connections to a given location. To account for biological variability and preparation-related differences in absolute latency, we report latency across the population as time elapsed from the first frame time, determined as described above. To generate average latency contour plots for age and hemisphere groups, contour plots were first calculated for individual movies, smoothed with a 5-pixel sliding box average across only the horizontal dimension (to aid alignment over the anterior-posterior axis), and averaged (Fig 1h).

For intracellular recording of spontaneous miniature inhibitory postsynaptic currents, excitatory neurons located in layer 4, approximately 350-450 um in depth from the cortical surface and 50–80 μm below the cut surface of the slice, were visualized using infrared gradient contrast optics and patched with glass electrodes (6–7 MOhm) containing the following intracellular solution (in mM) 128 K-methylsulfate, 4 MgCl, 10 HEPES, 1 EGTA, 4 NaATP, 0.4 NaGTP, 10 Na-phosphocreatine, and 0.01 QX-314 (pH 7.25 and 300 mOsm). The ACSF bath contained 1uM TTX and 50uM D-AP5. Recordings were made in whole-cell voltage clamp mode using a Multiclamp 700 A amplifier (Axon Instruments, Molecular Devices, Sunnyvale, California, USA). We measured inhibitory currents at a holding potential of 0 mV. We used the custom software package ephus (http://www.ephus.org) for instrument control and acquisition written in Matlab (MathWorks, Natick, MA, USA).

In the analyses reported for in vitro experiments, we used both custom software written in Igor Pro 9.2 (Wavemetrics, Lake Oswego, OR, USA) and MatLab R2021b (MathWorks, Natick, MA, USA). For statistical tests, we used Graphpad Prism version 9.5.1 (GraphPad Software, San Diego, California USA). All software implementation was executed in 64-bit Windows.

For statistical analyses across both VSD and mIPSC metrics, we first tested whether each group of measurements in the population was determined to be normally distributed by the Shapiro-Wilk test. When all groups were determined to be normally distributed, we performed Welch’s t-tests for multiple comparisons using the Holm-Šídák correction method. Otherwise, we used the non-parametric Dunn’s multiple comparisons test with Bonferroni correction. Within hemispheres, planned comparisons were made only between the youngest group and all other groups. Within group outliers were removed as determined by the MatLab function *isoutlier*, which identifies values more than 3 scaled median absolute deviations (MAD) away from the median value. Statistical analysis results and population source data will be made available via figshare. Raw data and other ancillary code are available upon reasonable request.

In vivo extracellular electrophysiological recordings were performed in mice aged P33-P56. We administered 75 mg/kg ketamine and 0.5 mg/kg medetomidine for anesthetized recordings. Anesthesia was supplemented during surgery and throughout the recordings as needed. Following anesthesia, mice were kept on a heating pad at 36-38°C and placed in a stereotaxic instrument equipped with head-fixed orbital bars, and a bite bar. We made a craniotomy (2x2mm^2^) and durotomy over the ACx, centered around 1.5mm anterior and 4mm lateral to lambda. The exposed cortex was kept moist with cortex buffer ((in mM) 125 NaCl, 5 KCl, 10 Glucose, 10 HEPES, 2 CaCl2, 2 MgSO4) throughout the recording session. A 32 channel, 2-shank silicone probe (P1, Cambridge Neurotech) was inserted into the auditory cortex at a depth of 0.6mm ± (0.1mm) from the tip of the probe. The probe’s recording sites spanned 250um, covering mainly the granular layer, but also supra- and/or sub-granular layers. The probe was lowered at a speed of ∼100um every 5 minutes. Recordings were obtained using Cheetah software (Neuralynx), with all channels sampled in continuous mode at 30.3kHz. All recordings were done in a sound-attenuated chamber, using a custom-built real-time Linux system (200kHz sampling rate) driving a Lynx-22 audio card (Lynx Studio Technology, Newport Beach, California, USA) with an ED1 electrostatic speaker (Tucker-Davis Technologies, Alachua, Florida, USA) in a free-field configuration (speaker located 6 inches lateral to, and facing the contralateral ear). The stimuli were created with custom MATLAB scripts to compute tuning curves. We used a set of pure tones (16 frequencies, 3 amplitudes - 20, 40, 60dB) that lasted 100ms, with an inter-stimulus interval of 1s.

For analysis of *in vivo* extracellular neuronal activity, we used Kilosort to extract spike times and determine putative spike clusters, followed by custom routines in MatLab to determine the tone response properties of identified spike clusters. A cluster was first determined to be responsive to tone presentations and included in further analysis based on the optimal kernel bandwidth of the peri-stimulus time histogram approach described by Shimazaki and Shinomoto 2010^35^. Tone frequency tuning curves for each tone responsive cluster were then computed by counting the total number of spikes occurring for each frequency in the 3 contiguous peristimulus time histogram (PSTH) bins with the highest spike counts. For each recording location, a population tuning curve was computed by averaging the normalized tuning curves of all criteria-passing spike clusters at that location. For each location, a tone frequency was considered to be within the response field if the average tuning curve at that frequency exceeded ⅔ of the maximum observed firing rate. To determine whether the proportion of 7kHz responsive recording locations was influenced by tone rearing in each hemisphere, each location was first classified as 7kHz responsive or not based on whether 7kHz +/- ⅓ octave tones were contained within the response field as described above. Counts were made for each contingency category as reported in Fig 3 and tested for significance using Fisher’s Exact Test.

