## Supplementary table 1 for "Mouse Auditory Cortex Undergoes Asynchronous Maturation in the Right and Left Hemispheres"

Figure 1f

| **Cortical depth 25th percentile** | **p-values**, Welch's ANOVA test, post hoc Dunnett's t test |
|---|---|
| LH <P14 vs. LH P14-P16 | ns |
| LH <P14 vs. LH >P16 | <0.0001 |
| RH <P14 vs. RH P14-P16 | 0.0214 |
| RH <P14 vs. RH >P16 | 0.0023 |
| **Cortical depth 50th percentile** | |
| LH <P14 vs. LH P14-P16 | ns |
| LH <P14 vs. LH >P16 | <0.0001 |
| RH <P14 vs. RH P14-P16 | 0.0022 |
| RH <P14 vs. RH >P16 | 0.0001 |
| **Cortical depth 75th percentile** | |
| LH <P14 vs. LH P14-P16 | ns |
| LH <P14 vs. LH >P16 | <0.0001 |
| RH <P14 vs. RH P14-P16 | 0.0025 |
| RH <P14 vs. RH >P16 | 0.0011 |

Figure 1g

| **VSD center of mass** | **p-values**, Welch's ANOVA test, post hoc Dunnett's t test |
|---|---|
| LH <P14 vs. LH P14-P16 | ns |
| LH <P14 vs. LH >P16 | <0.0001 |
| RH <P14 vs. RH P14-P16 | 0.0028 |
| RH <P14 vs. RH >P16 | 0.0001 |

Within hemisphere statistical data for Figure 1: Hemispheric differences in the maturation of thalamocortical input to the ACx.
